# The rice NLR pair Pikp-1/Pikp-2 initiates cell death through receptor cooperation rather than negative regulation

**DOI:** 10.1101/2020.06.20.162834

**Authors:** Rafał Zdrzałek, Sophien Kamoun, Ryohei Terauchi, Hiromasa Saitoh, Mark J Banfield

**Affiliations:** Department of Biological Chemistry, John Innes Centre, Norwich Research Park, Norwich, NR4 7UH, UK; The Sainsbury Laboratory, University of East Anglia, Norwich Research Park, Norwich, NR4 7UH, UK; Division of Genomics and Breeding, Iwate Biotechnology Research Centre, Iwate, Japan; Laboratory of Crop Evolution, Graduate School of Agriculture, Kyoto University, Kyoto, Japan; Laboratory of Plant Symbiotic and Parasitic Microbes, Department of Molecular Microbiology, Faculty of Life Sciences, Tokyo University of Agriculture, Tokyo 156-8502, Japan

## Abstract

Plant NLR immune receptors are multidomain proteins that can function as specialized sensor/helper pairs. Paired NLR immune receptors are generally thought to function via negative regulation, where one NLR represses the activity of the second and detection of pathogen effectors relieves this repression to initiate immunity. However, whether this mechanism is common to all NLR pairs is not known. Here, we show that the rice NLR pair Pikp-1/Pikp-2, which confers resistance to strains of the blast pathogen *Magnaporthe oryzae* (syn. *Pyricularia oryzae*) expressing the AVR-PikD effector, functions via receptor cooperation, with effector-triggered activation requiring both NLRs to trigger the immune response. To investigate the mechanism of Pikp-1/Pikp-2 activation, we expressed truncated variants of these proteins, and made mutations in previously identified NLR sequence motifs. We found that any domain truncation, in either Pikp-1 or Pikp-2, prevented cell death in the presence of AVR-PikD, revealing that all domains are required for activity. Further, expression of individual Pikp-1 or Pikp-2 domains did not result in cell death. Mutations in the conserved P-loop and MHD sequence motifs in both Pikp-1 and Pikp-2 prevented cell death activation, demonstrating that these motifs are required for the function of the two partner NLRs. Finally, we showed that Pikp-1 and Pikp-2 associate to form homo- and hetero-complexes in planta in the absence of AVR-PikD; on co-expression the effector binds to Pikp-1 generating a tripartite complex. Taken together, we provide evidence that Pikp-1 and Pikp-2 form a fine-tuned system that is activated by AVR-PikD via receptor cooperation rather than negative regulation.

## Introduction

Like animals, plants are constantly threatened by pathogens and pests. To defend themselves, they have evolved a sophisticated immune system that relies on both cell surface and intracellular receptors (1, 2). The majority of cloned resistance genes are intracellular immune receptors that belong to the nucleotide-binding, leucine-rich repeat (NLR) superfamily (3). NLRs activate immunity leading to disease resistance following recognition of pathogen elicitors, typically effectors delivered into host cells to promote pathogenesis (4). NLR-mediated immunity can include localised cell death known as the Hypersensitive Response (HR) (5), which contributes to limiting pathogen spread through host tissue.

The canonical architecture of plant NLRs consists of an N-terminal Toll/Interleukin-1 receptor homology (TIR) domain or coiled-coil (CC) domain ((including the RPW8-like CC, CC_R_), establishing the TIR-NLR, CC-NLR and CC_R_-NLR families), a central NB-ARC domain (Nucleotide-binding adaptor and APAF-1, R proteins, and CED-4), and a C-terminal leucine-rich repeat (LRR) domain. Conceptual frameworks for the roles of each domain are established, although their precise role may vary from one NLR to another (6). In brief, the N-terminal TIR or CC domains are thought to be involved in triggering cell death following effector perception, with recent studies suggesting a nucleotide hydrolase activity (for TIRs (7–9)) and membrane-perturbation (for oligomeric CCs (10, 11)). The NB-ARC domain acts as a molecular switch with the conformation of the protein stabilised by the bound nucleotide, ADP or ATP (12–15). Within the NB-ARC domain, several well-conserved sequence motifs are known, with the “P-loop” and “MHD” motifs located to the nucleotide binding site (16, 17). Mutations in these motifs have diverse effects on NLR activity. For example, mutations within the P-loop motif impair nucleotide binding, and often result in loss of protein function (18–20). Mutations in this motif can also prevent self-association and affect localisation (21). Mutations within the MHD motif frequently lead to constitutive activity (often called auto-activation (22–26)). The C-terminal LRR domain has a role in auto-inhibition (27–29), a function shared with animal NLRs (30–32), but can also define effector recognition specificity (33).

NLRs can function as singletons, capable of both perceiving effectors and executing a response (34, 35). This activity may require non-NLR interactors (36–40) or oligomerisation (41, 42). However, many NLRs require a second NLR for function, with three major classes described (43, 44). In each class, one of the NLRs functions as a “sensor” to detect the presence of the effector, whereas the second acts as a “helper”, and is required for cell death activity. For genetically linked sensor-helper NLR pairs, expression is driven from a shared promoter, and both proteins are required for effector perception (45). Interestingly, in many genetically linked NLR pairs, the sensor NLR contains an additional integrated domain that directly binds a pathogen effector (46–50). Integrated domains in NLRs have been found across all flowering plants (51–53). The separation of sensor/helper functions within NLR pairs may have evolutionary advantages, for example increasing tolerance to point mutations in the sensor (54).

CC-NLRs RGA5 and RGA4 from rice, and TIR-NLRs RRS1 and RPS4 from *Arabidopsis* are well established models in the study of genetically linked NLR pairs (45, 48, 55). RGA5 and RRS1 are the sensor NLRs (harbouring an integrated HMA (Heavy Metal Associated) domain and integrated WRKY domain respectively), and RGA4 and RPS4 are the helpers. In both systems, the helper NLRs appear to be auto-active when expressed alone in heterologous expression systems, and this auto-activity is suppressed on co-expression with the sensor NLR. Effector perception relieves suppression and initiates receptor activity (45, 56).

In rice, the CC-NLR pair Pik-1 and Pik-2 confers resistance to *Magnaporthe oryzae* (syn. *Pyricularia oryzae*) carrying the AVR-Pik effector. Similar to RGA5, Pik-1 has an integrated HMA domain, but unlike RGA5 this is positioned between the CC and NB-ARC domain, rather than after the LRR. The Pik-1 integrated HMA domain directly binds the AVR-Pik effector (50). However, how recognition of the effector translates into an immune response in the context of full-length receptors is unclear, as is the nature of any pre-activation state of the Pik-1/Pik-2 proteins. Further, which NLR domains are necessary and sufficient for immune signalling in this pair is unknown.

We previously showed that the AVR-Pik elicited hypersensitive cell death mediated by the Pik NLR pair can be recapitulated using transient expression in leaves of the model plant *Nicotiana benthamiana* (50, 57, 58). In this study, we investigated the roles and requirements of domains in the Pik NLR alleles Pikp-1 and Pikp-2 in planta using the *N. benthamiana* experimental system. We show that intact, full-length, Pikp-1 and Pikp-2 are necessary for a cell death response upon effector perception. Truncation of any domain results in lack of effector-dependent cell death compared to wild-type. Further, expression of any specific NLR domain, or combination of domains, does not result in cell death. We also show that native P-loop and MHD-like motifs are required in both Pikp-1 and Pikp-2 proteins for receptor activity. Finally, we demonstrate that Pikp-1 and Pikp-2 are able to form homo- and hetero-complexes in planta in the absence of the AVR-PikD. Upon binding of the AVR-PikD effector, a tri-partite complex is formed that may represent the activated state of the receptor.

## Materials and Methods

### Cloning

Domesticated sequences of full-length Pikp-1 and Pikp-2 (as described in (58)), and MLA10, were assembled into the pICH47751 vector under the control of the *mas* promoter and with C-terminal epitope tags (3x FLAG tag, V-5 tag or 6xHA tag accordingly) using the Golden Gate system (59). To obtain Pikp-1 and Pikp-2 individual domains and truncation variants, relevant sequences were amplified by PCR using the plasmids above as templates, and assembled into the pICH47751 vector under control of CaMV35S promoter and with C-terminal epitope tags (6xHis + 3xFlag (HellFire (HF)) for Pikp-1 derivatives and 6xHA for Pikp-2 derivatives) using the Golden Gate system. Myc:AVR-PikD and Myc:AVR-PikD^H46E^ constructs used were as described in (58). All DNA constructs were confirmed by sequencing and transformed into *Agrobacterium tumefaciens* strain GV3101 via electroporation.

### Mutagenesis

To generate Pikp mutants (P-loop and MHD-like motifs), we introduced mutations into the relevant NB-ARC domain modules using site-directed mutagenesis. Subsequently these domain constructs were used to generate full length NLRs by assembly using the Golden Gate system. Each of the constructs were assembled with the CaMV35S promoter with relevant tags (6xHis + 3xFlag (HellFire (HF)) for Pikp-1 derivatives and 6xHA tag for Pikp-2 derivatives).

### Cell death assays

*Agrobacterium tumefaciens* strain GV3101 carrying the appropriate constructs were suspended in infiltration buffer (10 mM MgCl2, 10 mM MES, pH 5.6, 150 mM acetosyringone) and mixed prior to infiltration at the following final OD600: NLRs and NLR-derivatives 0.4, effectors 0.6, P19 (silencing suppressor) 0.1. Bacteria were infiltrated into leaves of ~4 weeks old *N. benthamiana* plants using a 1ml needleless syringe. At 5 days post infiltration (dpi), detached leaves were imaged under UV light on the abaxial side, and visually scored for cell death response (see below). To confirm protein expression, representative infiltration spots were prepared, frozen in liquid nitrogen, ground to a fine powder, mixed with extraction buffer (see below) in 2 ml/g ratio, centrifuged, mixed with loading dye and loaded on an SDS-PAGE gel for western blot analysis.

### Cell death scoring

Pictures of the leaves at 5 dpi were taken as described previously (57) and cell death (visible as green fluorescence area under the UV light) was scored according to the scale presented in (50). The dot plots were generated using R v3.4.3 (https://www.r-project.org/) and the graphic package ggplot2 (60). Dots represent the individual datapoints and the size of larger circles is proportional to the number of dots within that score. Dots of the same colour within one plot come from the same biological repeat. All positive and negative controls were also scored and are represented on relevant plots. As positive and negative controls were included on most leaves there are more data points for these samples.

### Co-Immunoprecipitation

Protein extraction was conducted as described in (61) with minor modifications. Extraction buffer GTEN (10% glycerol, 25 mM Tris, pH 7.5, 1 mM EDTA, 150 mM NaCl), 2% w/v PVPP, 10 mM DTT, 1× protease inhibitor cocktail (Sigma), 0.1% Tween 20 (Sigma) was added to frozen tissue in 2 ml/g ratio. The sample was resuspended and centrifuged for 30 min (4500g) at 4°C. The supernatant was filtered through a 0.45 μm filter. Anti-FLAG M2 magnetic beads (Sigma, M8823) were washed with the IP buffer (GTEN + 0.1% Tween 20), resuspended, and added to protein extracts (20 μl of resin per 1.5 ml of extract). Samples were incubated for an hour at 4°C with gentle shaking. Following incubation, the resin was separated using magnetic stand and washed 5 times with IP buffer. For elution, beads were mixed with 30 μl of Loading Dye and incubated at 70°C for 10 min. Finally, samples were centrifuged and loaded on a precast gradient gel (4-20%, Expedeon) for western blot analysis.

### Western blot

Western blots were performed as described previously (61). Following SDS-PAGE, proteins were transferred onto PVDF membrane using Trans-Blot Turbo Transfer Kit (Biorad) and blocked in 5% milk in TBS-T (50mM Tris-HCl, 150mM NaCl, 0.1% Tween20, pH 8.0) at 4°C for at least 1 hour. Respective primary HRP-conjugated antibodies (α-FLAG: Cohesion Biosciences, CPA9020; α-HA: Invitrogen, #26183-HRP; α-V-5: Invitrogen, #MA5-15253-HRP; α-Myc (9E10): Santa Cruz Biotechnology, SC-40) were applied for overnight incubation (4°C). Membranes were then rinsed with TBS-T. Proteins were detected using ECL Extreme reagents (Expedeon) in chemiluminescence CCD camera (ImageQuant LAS 500).

## Results

### Each domain of Pikp-1 and Pikp-2 is required for receptor activation

To investigate the roles and requirements for individual domains of Pikp-1 (CC, HMA, NB-ARC and LRR) and Pikp-2 (CC, NB-ARC and LRR) in triggering cell death, we transiently expressed each of these in *N. benthamiana* using *Agrobacterium tumefaciens* mediated transformation (henceforth agroinfiltration). All constructs were tagged at their C-terminus with the HellFire tag (6xHis + 3xFlag (HF), for Pikp-1 domains) or HA tag (for Pikp-2 domains). The boundaries of the domains used were as defined in (50).

We found that each of the individual domains of either Pikp-1 or Pikp-2 were unable to trigger cell death when expressed alone, or in the presence of the corresponding paired NLR and/or effector (Fig 1, Fig 2, S1 Fig). We confirmed that all the proteins accumulated to detectable levels using western blot analysis (S2 Fig). We then systematically truncated Pikp-1 or Pikp-2 at relevant domain boundaries, and expressed these alone or in the presence of the corresponding paired NLR and/or effector, to search for any minimum functional unit (Fig 1B, Fig 2A, Fig 2B, S1 Fig). In all cases tested no cell death was observed, despite the proteins accumulating in plant tissues (S2 Fig). The only combination that gave cell death was the positive control of full length Pikp-1 and Pikp-2 in the presence of AVR-PikD. These results show that the Pikp-1/Pikp-2 pair work together to deliver a cell death response on effector perception, and all domains are required for activity.

**Fig 1.**
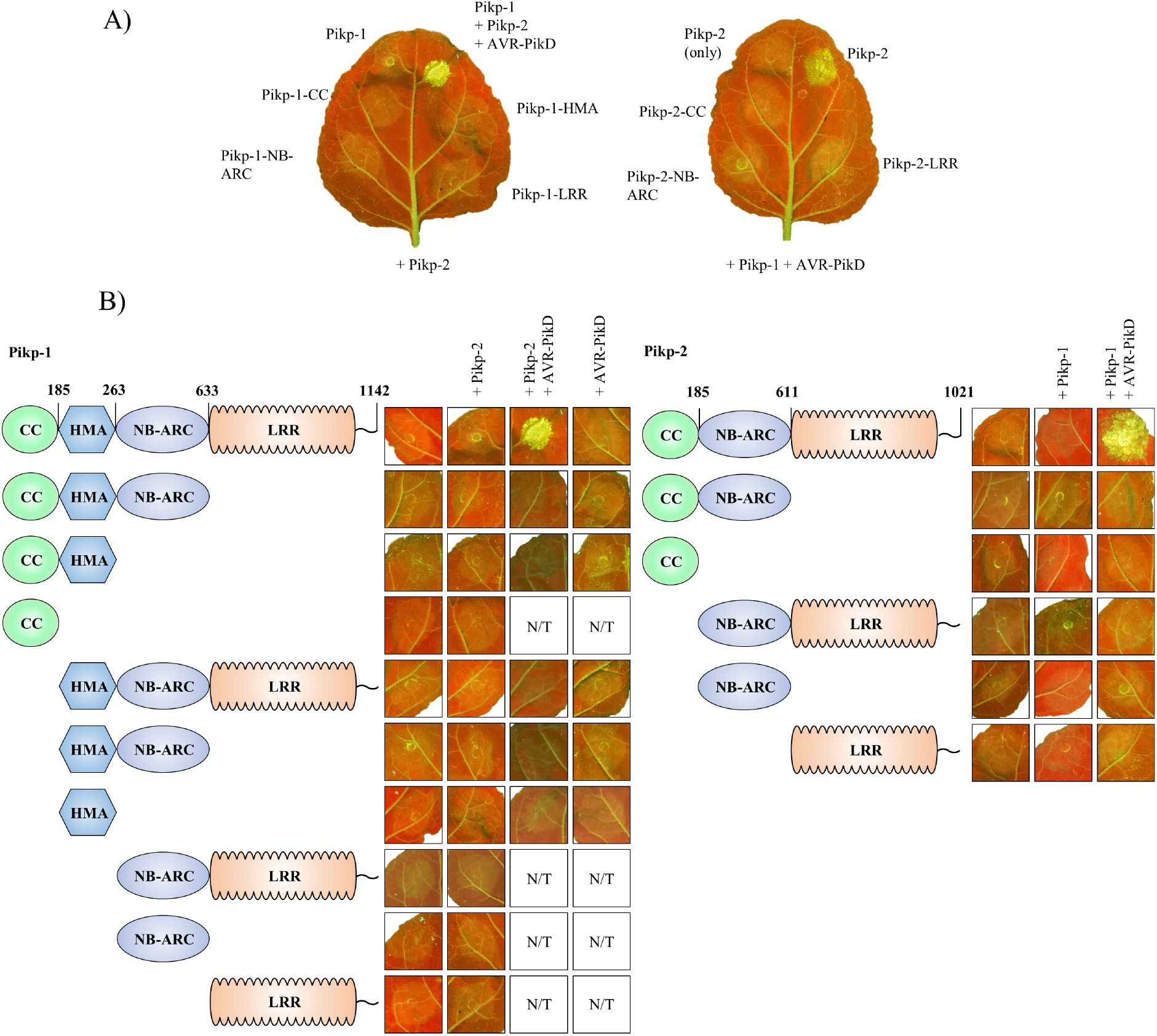
Each domain of Pikp-1 and Pikp-2 is required for receptor activation. **A)** Representative *N. benthamiana* leaves showing that expression of the individual domains of Pikp-1 (left) or Pikp-2 (right) were unable to elicit a cell death response in presence of the corresponding paired NLR (for Pikp-1) or paired NLR and effector (for Pikp-2). Pikp-1+Pikp-2+AVR-PikD is shown as a positive control. **B)** Representative agroinfiltration spots show that the truncated variants of Pikp-1 (left) or Pikp-2 (right) were unable to elicit a cell death response, either when overexpressed alone, or in the presence of corresponding full-length NLR and/or effector. Combinations of constructs without HMA domain were not tested (N/T) in presence of the effector.

**Fig 2.**
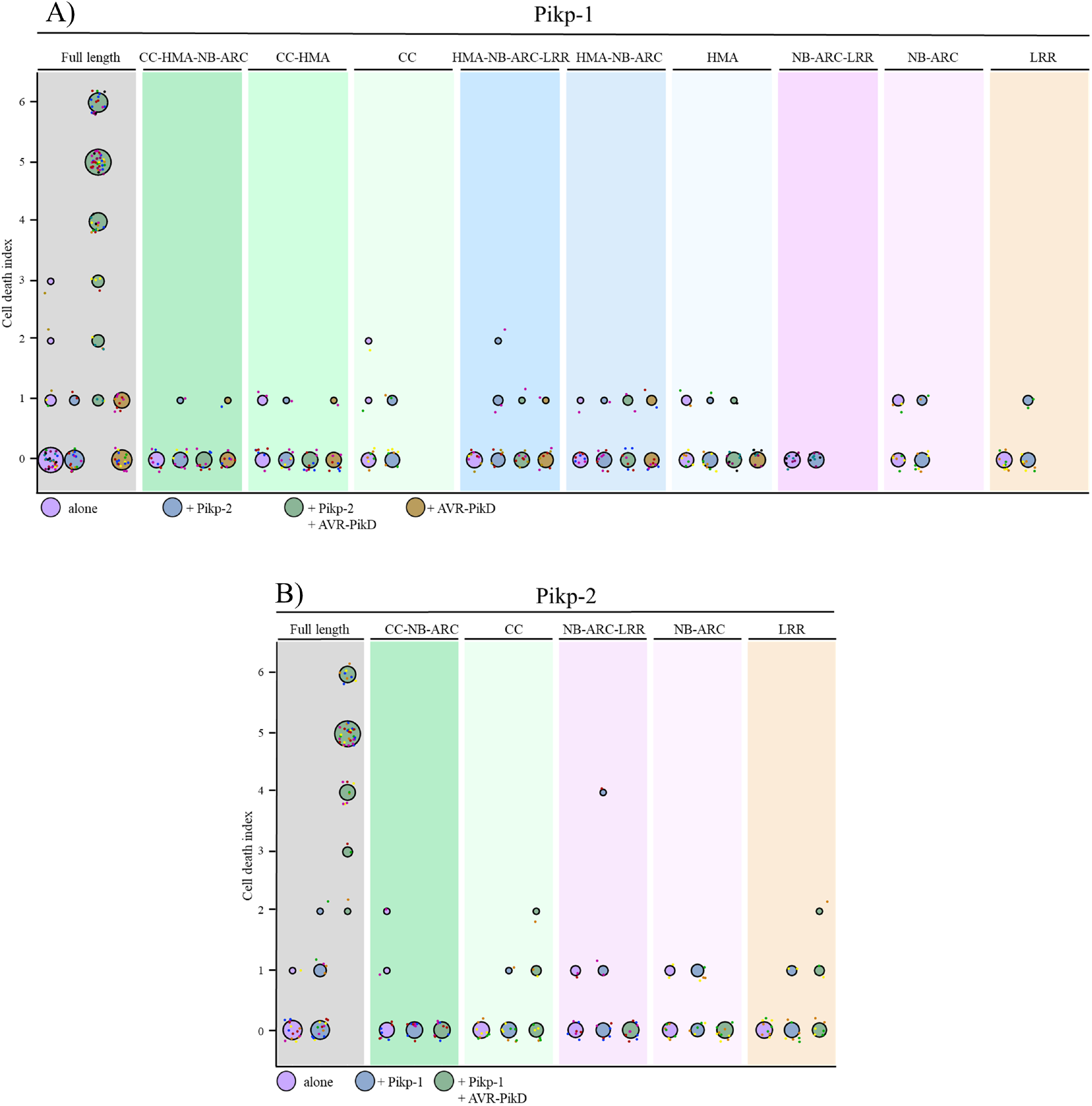
Each domain of Pikp-1 and Pikp-2 is required for receptor activation. Cell death quantification for the infiltration combinations of Fig 1 shown as dot plots, for Pikp-1 **(A)** and Pikp-2 **(B)** respectively. Each of the dots has a distinct colour corresponding to the biological replicate, and are plotted around the cell death score for visualization purposes. Each set of infiltrations were repeated in 3 biological replicates with at least 2-3 technical replicates. The size of the central dot at each cell death value is proportional to the number of replicates of the sample with that score.

### Conserved NB-ARC domain sequence motifs are required for Pikp-1 and Pikp-2 activity

Next, we tested whether previously characterised sequence motifs within the nucleotide-binding pocket of the Pikp-1 and Pikp-2 NB-ARC domains are required for receptor activity. Firstly, we generated mutations in the P-loop motifs of Pikp-1 and Pikp-2 (Pikp-1^K296R^ and Pikp-2^K217R^). Such mutations restrict nucleotide binding, and have previously been shown to impair NLR function (18, 19, 62, 63). On expression in *N. benthamiana* via agroinfiltration, we found that these mutations abolish cell death activity in planta, including when expressed in the presence of the paired NLR and the AVR-PikD effector (Fig 3, S3A Fig). This reveals that an intact P-loop motif is required in both Pikp-1 and Pikp-2 for activity. Expression of all proteins was confirmed by western blot analysis (S3B Fig). Secondly, we generated mutations in the “MHD” motifs of Pikp-1 and Pikp-2. Although classically defined as Methionine-Histidine-Aspartate (MHD), the residues that comprise this motif in plant NLRs can vary. Here we will refer to this as the MHD-like motif. In Pikp-1, the MHD-like motif residues are Ile-His-Pro (IHP), while in Pikp-2 they are Val-His-Asp (VHD). Mutations within this NLR motif frequently lead to auto-activation and cell death in the absence of pathogen perception (64–66). To determine the importance of the MHD-like motif for Pikp-1 and Pikp-2 activity, we generated triple alanine mutants of each protein (Pikp-1^599IHP601→AAA^, and Pikp-2^557VHD559→AAA^) and a Pikp-2^D559V^ mutant. On expression in *N. benthamiana* via agroinfiltration, we found that expression of these mutants alone did not result in auto-activity and cell death (Fig 3). We also found that any combination of the MHD-like motif mutants with wild-type or mutant paired NLRs, with or without the AVR-PikD effector, did not result in cell death (Fig 3B, Fig 3D). These results show that the native MHD-like motifs of Pikp-1 and Pikp-2 are required to trigger cell death. All proteins were expressed to detectable levels, as confirmed by western blot analysis (S3B Fig).

**Fig 3.**
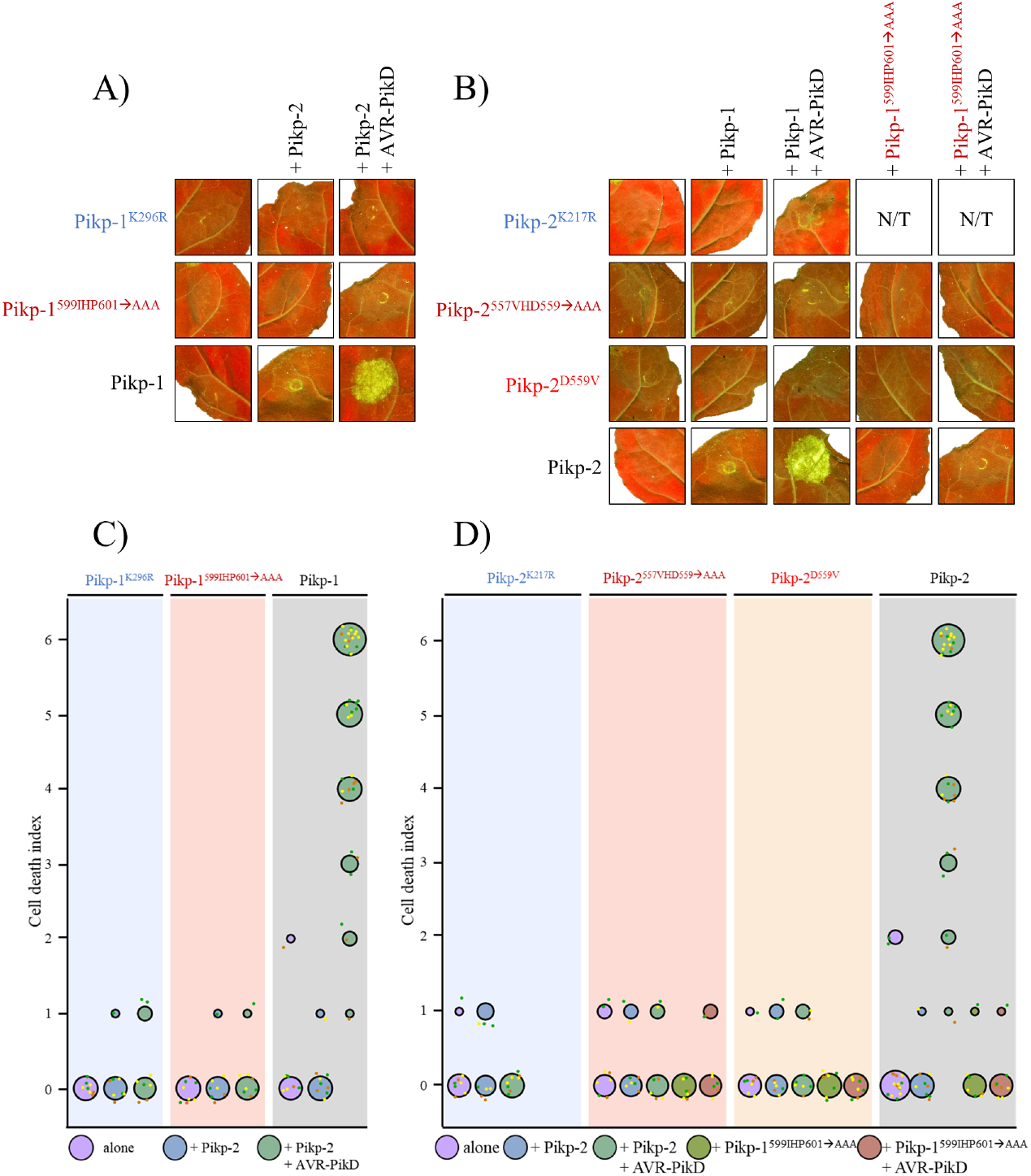
Conserved NB-ARC domain sequence motifs are required for Pikp-1 and Pikp-2 activity. **A)** Mutation of the P-loop motif of Pikp-1 (Pikp-1^K296R^) results in loss of cell death response upon effector perception. Mutation of the Pikp-1 MHD-like motif (Pikp-1^599IHP601→AAA^) does not lead to auto-activity when overexpressed alone, or in the presence of corresponding intact NLR. Further, this mutant was also unable to trigger a cell death response when co-expressed with AVR-PikD. **B)** Mutation of the P-loop motif of Pikp-2 (Pikp-2^K217R^) results in loss of cell death response upon effector perception. Mutation of the Pikp-2 MHD-like motif (Pikp-2^557VHD559→AAA^ and Pikp-2^D559V^) does not lead to auto-activity when overexpressed alone, in the presence of corresponding intact NLR, or its MHD-like mutant. Further, these mutants were also unable to trigger a cell death response when co-expressed with AVR-PikD. Each set of infiltrations were repeated in 3 biological replicates with at least 2-3 technical replicates within each. The square showing the infiltration spot for wild type Pikp-1+Pikp-2 was as-used in Fig. 1B. Squares representing Pikp-1^599IHP601àAAA^+Pikp-2 and Pikp-1^599IHP601àAAA^+Pikp-2+AVR-PikD are the same on both panels, presented for comparison. **C)** and **D)** Cell death quantification for each infiltration shown as dot plots. Each of the dots has a distinct colour corresponding to the biological replicate, and are plotted around the cell death score for visualization purposes. The size of the central dot at each cell death value is proportional to the number of replicates of the sample with that score. The data for Pikp-1+Pikp-2 (wild type) is a subset of the previous experiment (Fig 2A, Fig 2B), used here for comparison. The data shown for Pikp-1+Pikp-2, Pikp-1+Pikp-2+AVR-PikD, Pikp-1^599IHP601^+Pikp-2 and Pikp-1^599IHP601^+Pikp-2+AVR-PikD are the same in both panels, presented for comparison.

### Pikp-1 and Pikp-2 form homo- and hetero-complexes in planta

Paired NLRs can form homo- and hetero-complexes in planta (45, 48). To investigate whether Pikp-1 and Pikp-2 can also homo- and/or hetero-associate, both in the absence and in the presence of the effector, we performed in planta co-immunoprecipitation (co-IP) assays. To test for homo-complex formation we expressed differentially tagged Pikp-1 constructs (FLAG tag and V-5 tag), or Pikp-2 constructs (FLAG tag and HA tag) in *N. benthamiana* via agroinfiltration, followed by immunoprecipitation with α-FLAG resin. The barley NLR MLA10 (expressed with a FLAG tag) served as a negative control for interactions. Each FLAG-tagged protein was expressed, and immunoprecipitated as expected (lower panels, Fig 4A, Fig 4B). For Pikp-1, we observe co-immunoprecipitation of Pikp-1:V-5 with Pikp-1:FLAG, but not with MLA10:FLAG, and Pikp-1:V-5 did not show non-specific interaction with the resin when expressed alone (Fig 4A). Similar results were obtained for Pikp-2 (Fig 4B), where Pikp-2:HA immunoprecipitated Pikp-2:FLAG on co-expression. Faint bands of Pikp-2:HA were also observed with MLA10:FLAG. However, a similar band can be observed where Pikp-2:HA is expressed alone, indicating a weak non-specific binding to the resin. The presence of the AVR-PikD effector (or the mutant AVR-PikD^H46E^ as a negative control) does not affect the homo-association of Pikp-1 or Pikp-2 (S4 Fig).

**Fig 4.**
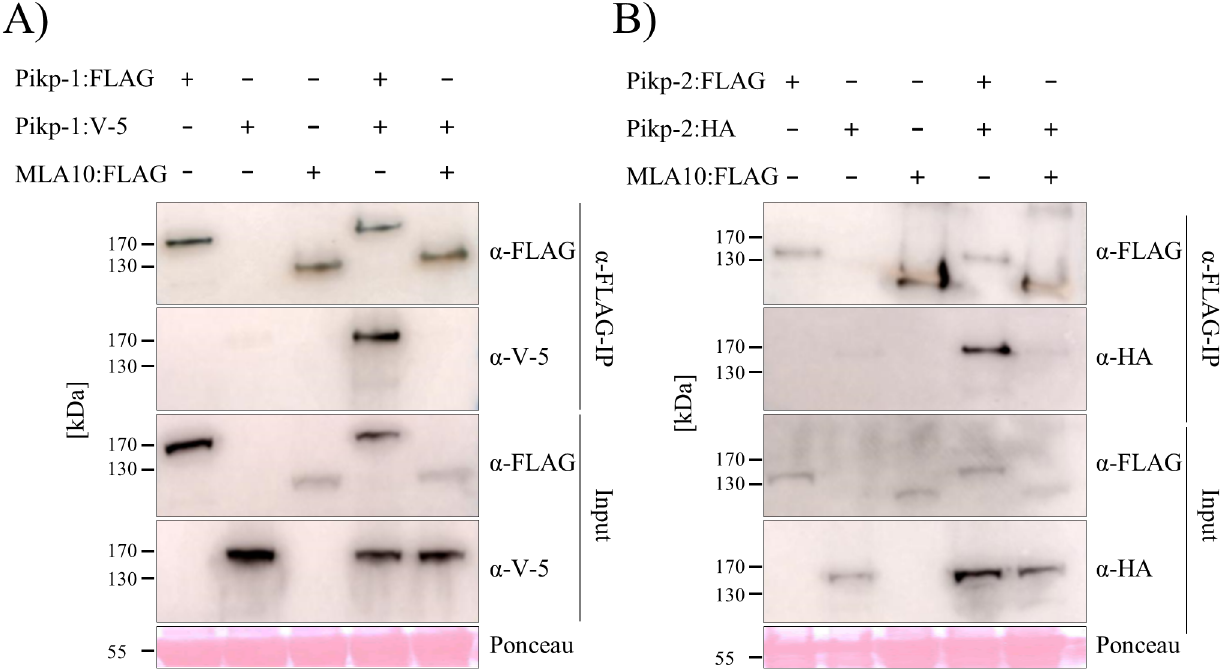
Pikp-1 and Pikp-2 form homo-complexes. **A)** Pikp-1:FLAG, Pikp-1:V-5 and MLA10:FLAG and **B)** Pikp-2:FLAG, Pikp-2:HA and MLA10:FLAG were expressed alone or in the combinations shown. Subsequently, anti-FLAG immunoprecipitation (α-FLAG-IP) was performed, followed by western blot analysis with relevant antibodies to detect the proteins (upper panel). The lower panel confirms presence of all the proteins prior to immunoprecipitation. Experiments were repeated at least 3 times with similar results.

We then tested whether Pikp-1 and Pikp-2 can form hetero-complexes. Using the resources described above, we co-expressed the proteins and performed α-FLAG pull downs. We show that Pikp-2:HA co-immunoprecipitated with Pikp-1:FLAG, but not with MLA10:FLAG, indicating that these NLRs specifically hetero-associate (Fig 5A). We also tested whether co-expression with AVR-PikD affects the formation of Pikp-1/Pikp-2 hetero-complexes. We co-expressed Pikp-1:V-5, Pikp-2:FLAG and Myc:AVR-PikD followed by α-FLAG pull down (note: in this case Pikp-2:FLAG is immunoprecipitated). All three proteins could be detected after α-FLAG pull down (Fig 5B). We suggest that Pikp-2 associates with Pikp-1, which is also bound to the AVR-PikD effector, forming tri-partite complex. Co-expression with the AVR-PikD^H46E^ mutant was used as a negative control for effector interaction.

**Fig 5.**
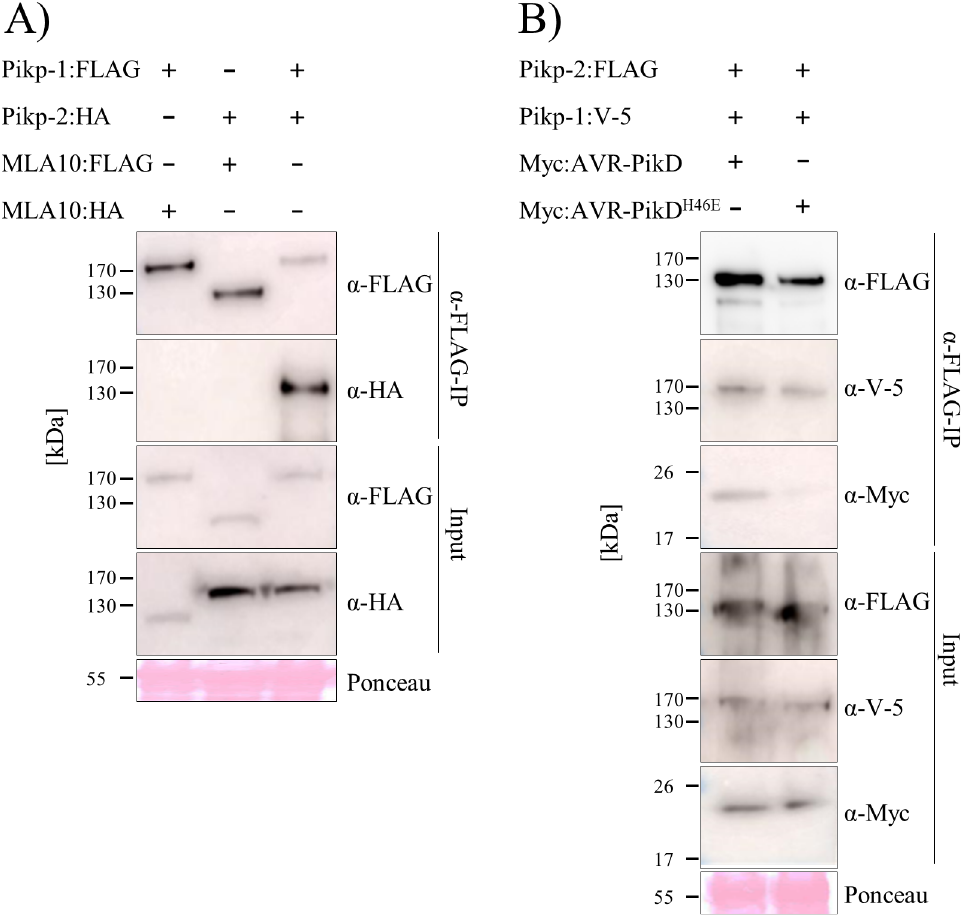
Pikp-1 and Pikp-2 form hetero-complexes prior to and upon recognition of AVR-PikD. **A)** Pikp-1:FLAG, Pikp-2:HA, MLA10:FLAG and MLA10:HA and **B)** Pikp-2:FLAG, Pikp-1:V-5, Myc:AVR-PikD and Myc:AVR-PikD^H46E^ were expressed in the combinations shown. Subsequently, anti-FLAG immunoprecipitation (α-FLAG-IP) was performed, followed by western blot analysis with relevant antibodies to detect the proteins (upper panel). The lower panel confirms presence of all the proteins prior to immunoprecipitation. Experiments were repeated at least 3 times with similar results.

## Discussion

Genetically linked NLR pairs are emerging as an important class of immune receptor in plants. Established models for paired NLR receptors suggest they function via negative regulation where a sensor NLR, that often carries an integrated domain, represses the activity of the second. Binding of pathogen effectors to the sensor NLR relieves this negative regulation. In this study, we show that the rice NLR pair Pikp-1/Pikp-2 differs from this model and works via receptor cooperation. Pikp-2 is not auto-active when expressed in the absence of Pikp-1. Both Pikp-1 and Pikp-2 are required to trigger cell death upon binding of the AVR-PikD effector to the integrated HMA domain of Pikp-1, and all the domains are indispensable for this activity. Further, we determined the requirements for conserved NB-ARC domain sequence motifs, the P-loop and MHD-like motifs. Finally, we find Pikp-1 and Pikp-2 can form homo- and hetero-complexes that are likely important for function.

The expression of individual domains of a number of NLRs can result in cell death. In particular, CC domains and other N-terminal truncations can induce cell death when expressed in planta (19, 42, 67–70). This is thought to reflect oligomerization of the CC domains, resulting in minimal functional units that can trigger cell death. However, the CC domains of either Pikp-1 or Pikp-2 did not display cell death inducing activity. This likely reflects an inability of these domains to adopt a configuration that supports cell death when expressed alone. Further, we did not observe cell death on expression of full-length Pikp-1 with the Pikp-2 CC domain (with or without AVR-PikD). We consider this a biologically relevant test for CC domain-mediated cell death in a paired NLR compared to co-expression of the CC domains with short epitope tags, or fused to GFP/YFP (a strategy required to observe cell death for some CC domains (38, 39, 71), but not used here). It is possible that further studies may identify a Pikp-1 or Pikp-2 CC domain construct that supports cell death, as studies with MLA10 family NLRs showed that a single amino acid change can make the difference between observing cell death or not (69), and chimeric NLRs with swaps within the CC domains can result in cell death (72).

Considering NLR regions other than the N-terminal domains, expression of the NB-ARC from Rx resulted in cell death (73). However, we did not observe this phenotype on expression of the NB-ARC domains of Pikp1 or Pikp-2. For the NLR RPS5, it was shown that a CC-NB-ARC construct can elicit cell death (40), but this may be due to deletion of the LRR domain that may have a role in auto-inhibition prior to effector detection (28). We did not observe cell death following deletion of the LRR domains of Pikp-1 or Pikp-2. Together, our data shows that full-length Pikp proteins, and perception of the effector, are required for cell death activity in planta. This is an effective strategy to prevent mis-regulation of receptor activity in the absence of the pathogen, but highlights the need for additional studies to understand the molecular mechanistic basis of Pikp activation, and the diversity of paired NLR function more generally.

Although the P-loop motif is required for NLRs reported to work as singletons (18, 20), it is not always necessary for paired and networked NLRs (45, 56, 74). In genetically linked pairs RGA5/RGA4 and RRS1/RPS4, the helper NLR requires an intact P-loop for cell death, but not the sensor (45, 56). In CC_R_-type helper NLRs, such as ADR1 and NRG1 (which function downstream of several other NLRs, but are not genetically linked (75)), an intact P-loop motif may not be required (76, 77). Pikp-1 and Pikp-2 appear to function similar to the NRC network of solanaceous plants, where both sensor and helper NLRs require a native P-loop motif for function (74). So why do Pikp-1 and Pikp-2 both require a native P-loop? It maybe this just provides an additional layer of regulation. It is also possible that mutations in the P-loop affect protein folding by preventing ADP binding, and Pikp-1 is more sensitive to this than other sensor NLRs studied, or that ADP/ATP exchange is more important for transducing effector binding by the HMA integrated domain in Pikp-1, possibly determined by the unusual position of the integrated domain between the CC and NB-ARC domain in this NLR.

Residues of the MHD-like motif are involved in binding ADP in the inactive state of NLRs (12, 78), and mutations in this motif can lead to auto-activity (22, 23). Mutations in the MHD-like motif of Pikp-1 and Pikp-2 are not auto-active, and result in a loss of cell death activity when expressed with the AVR-PikD effector. In RGA5 and RGA4, residues of the MHD-like motif are LHH and TYG, respectively, and the presence of a Glycine (G) in the third position of RGA4 was shown to be linked to RGA4 auto-activity, whereas introducing mutations into MHD-like motif of RGA5 did not abolish its ability to repress RGA4 (45). The most straightforward explanation for why changes at the MHD-like motif in Pikp-1 and Pikp-2 results in a loss of any cell death activity, rather than autoactivation, is that these mutations do not support the protein confirmation required, perhaps in the context of this NLR pair specifically.

Plant NLRs can form both homo- and hetero-complexes both prior to and after effector recognition (21, 40, 42, 79, 80), or undergo effector induced oligomerisation (81). In addition to the CC domains of CC-NLRs, the N-terminal TIR domains of TIR-NLRs have been shown to oligomerise (82–84). Recently, the structure of full-length ZAR1 revealed the role of oligomerisation in activation of a full-length NLR (11). Here, we have shown that Pikp-1 and Pikp-2 form both homo- and hetero-complexes in the absence and presence of the AVR-PikD effector. However, the conformation of the proteins, their stoichiometry, and their specific arrangement within the complexes, remain to be determined. Various models are possible for the active complex including a Pikp-1/Pikp-2 dimer, a higher order oligomer including multiple copies of the dimer, or a structure where Pikp-1 initiates the oligomerisation of Pikp-2 similar to the mechanism seen for NAIP2/NLRC4 (85) and NAIP5/NLRC4 (86).

In summary, our findings reveal that the Pikp-1/Pikp-2 NLR pair function via receptor cooperation rather than a suppression/activation mechanism, and signalling in planta requires the full-length proteins with native sequences at the P-loop and MHD-like sequence motifs. This suggests multiple mechanisms of regulation exist for NLRs. It is important to further investigate these mechanisms if we are to fully understand NLR function and use this to engineer improved disease resistance phenotypes in crops.

## Acknowledgements

This work was supported by the UKRI Biotechnology and Biological Sciences Research Council (BBSRC) Norwich Research Park Biosciences Doctoral Training Partnership, UK [grant BB/M011216/1]; the UKRI BBSRC, UK [grants BB/P012574, BB/M02198X]; the European Research Council [ERC; proposal 743165]; the John Innes Foundation; the Japan Society for the Promotion of Science (JSPS KAKENHI; proposal 18K05657, 15H05779, and 20H00421), and by the Strategic Research Project from Tokyo University of Agriculture. We thank Juan Carlos De la Concepcion and Josephine Maidment for help with cloning. We also thank Thorsten Langner and Hiroaki Adachi for comments on the manuscript.

