## Supplemental Data for "The rice NLR pair Pikp-1/Pikp-2 initiates cell death through receptor cooperation rather than negative regulation"

### Supplementary data

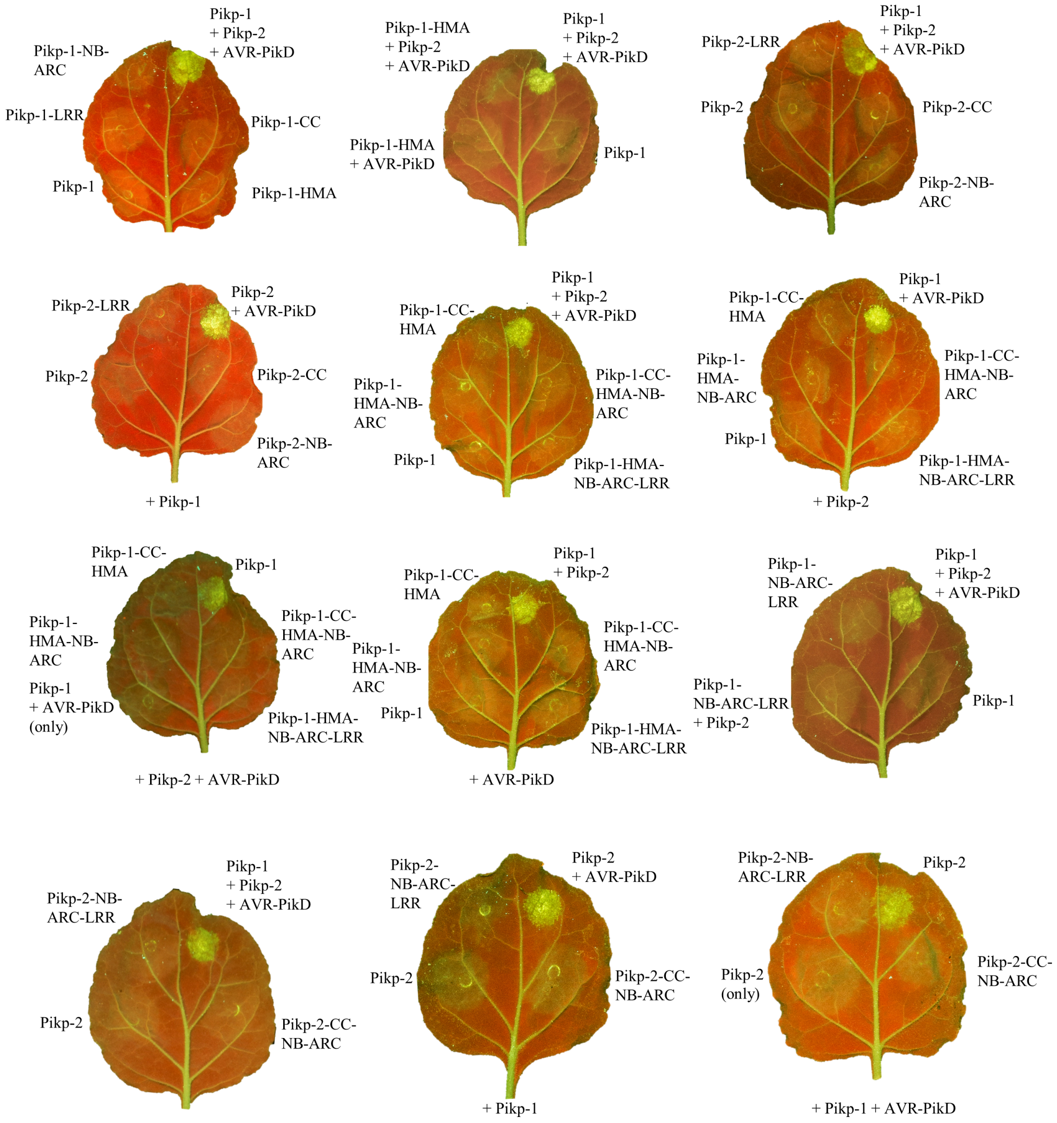


**S1 Fig. Each domain of Pikp-1 and Pikp-2 is required for receptor activation.** Representative *N. benthamiana* leaves showing that the individual domains and truncated variants of Pikp-1 or Pikp-2 were unable to elicit a cell death response, either when overexpressed alone, or in the presence of corresponding full-length NLR and/or effector. Individual infiltration spots from these leaves were used in Fig. 1B.

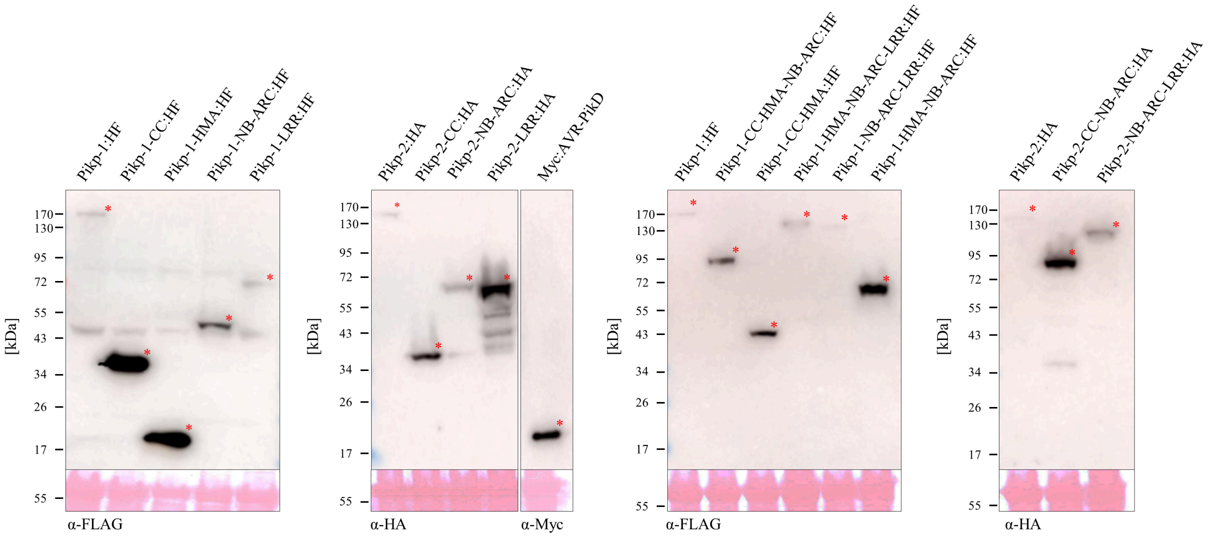


**S2 Fig. NLR and effector proteins were expressed to detectable levels in plant tissue.** Proteins were extracted from *N. benthamiana* leaf infiltrations (with individual constructs) and detected by western blot analysis with appropriate antibodies. Red asterisks indicate the bands of expected size for the proteins.

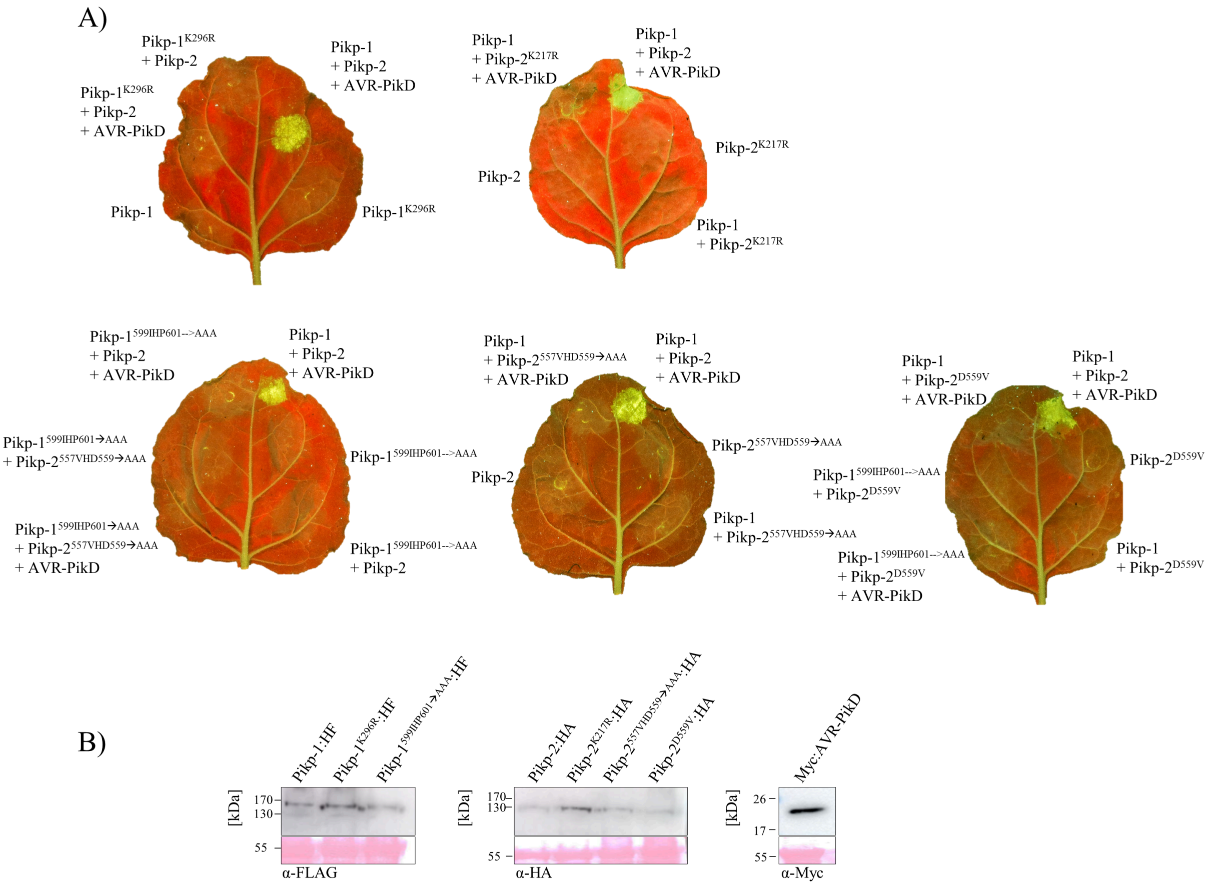


**S3 Fig. Conserved NB-ARC domain sequence motifs are required for Pikp-1 and Pikp-2 activity. A)** Representative *N. benthamiana* leaves showing that mutations in either of the P-loop or MHD-like motifs of Pikp-1 or Pikp-2 result in a loss of cell death response upon effector perception. Individual infiltration spots from these leaves were used in Fig. 2. **B)** Western blot analysis showing that proteins were expressed to detectable levels. Samples were taken from infiltrations with individual constructs.

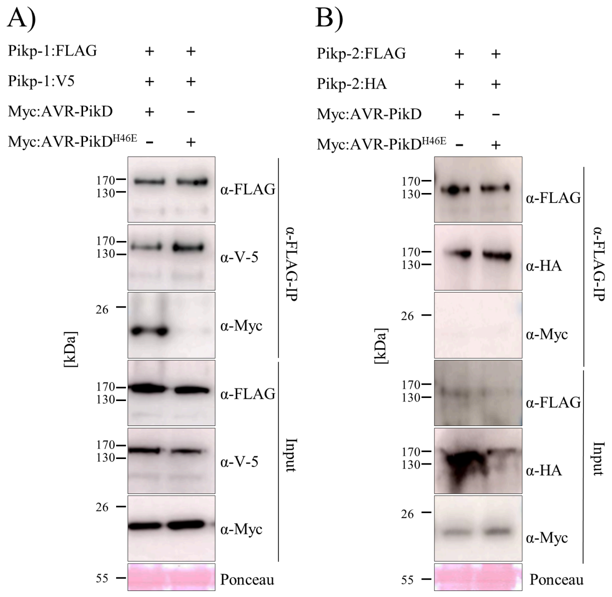


**S4 Fig. Presence of the AVR-PikD effector does not affect homo-association of Pikp-1 or Pikp-2**. **A)** Pikp-1:FLAG, Pikp-1:V-5, Myc:AVR-PikD and Myc:AVR-PikD$^{H46E}$ and **B)** Pikp-2:FLAG, Pikp-2:HA, Myc:AVR-PikD and Myc:AVR-PikD$^{H46E}$ were expressed in combinations shown. Subsequently, anti-FLAG immunoprecipitation (α-FLAG-IP) was performed, followed by western blot analysis with relevant antibodies to detect the proteins (upper panel). The lower panel confirms presence of all the proteins prior to immunoprecipitation. Experiments were repeated at least 3 times with similar results.
